# Does batrachotoxin autoresistance co-evolve with toxicity in *Phyllobates* poison-dart frogs?

**DOI:** 10.1101/460865

**Authors:** Roberto Márquez, Valeria Ramírez-Castañeda, Adolfo Amézquita

## Abstract

Toxicity is widespread among living organisms, and evolves as a multimodal phenotype. Part of this phenotype is the ability to avoid self-intoxication (autoresistance). Evolving toxin resistance can involve fitness tradeoffs, so autoresistance is often expected to evolve gradually and in tandem with toxicity, resulting in a correlation between the degrees of toxicity and autoresistance among toxic populations. We investigate this correlation in *Phyllobates* poison frogs, notorious for secreting batrachotoxin (BTX), a potent neurotoxin that targets sodium channels, using ancestral sequence reconstructions of BTX–sensing areas of the muscular voltage-gated sodium channel. Reconstructions suggest that BTX resistance arose at the root of *Phyllobates*, coinciding with the evolution of BTX secretion. After this event little or no further evolution of autoresistance seems to have occurred, despite large increases in toxicity throughout the history of these frogs. Our results therefore provide no evidence in favor of an evolutionary correlation between toxicity and autoresistance, which conflicts with previous work. Future research on the functional costs and benefits of mutations putatively involved in BTX resistance, as well as their prevalence in natural populations should shed light on the evolutionary mechanisms driving the relationship between toxicity and autoresistance in *Phyllobates* frogs.

---

A wide variety of species across the tree of life accumulate toxins as defenses from predators and parasites (Edmunds 1974; Mebs 2001). Toxicity usually evolves as a multi-level phenotype, comprised of physiological, behavioral, and morphological traits involved in acquiring, storing, delivering, and resisting toxins. The ability to avoid self-intoxication, also known as autoresistance, is an important piece of this phenotypic syndrome: For toxins to represent a selective advantage their bearer must not suffer their adverse effects. Predictably, toxic organisms display multiple auto–resistant phenotypes, such as specialized glands or organelles to compartmentalize toxins, or molecular changes in the toxins’ targets that inhibit or decrease their effects (Daly et al. 1980; Zhou and Fritz 1994; Geffeney et al. 2005; Zhen et al. 2012; Hanifin and Gilly 2015).

Although resistance can preexist toxicity, and therefore facilitate its evolution, evolving toxin resistance often involves functional changes that can have adverse pleiotropic effects, such as changes in nerve function (e.g. Brodie and Brodie 1999; Feldman et al. 2012) or reproductive output (e.g. Groeters et al. 1994; Gassmann et al. 2009). Therefore, autoresistance is usually thought to evolve gradually and in tandem with toxicity, with low levels of resistance allowing for gradual increases in toxicity that in turn promote small increases in resistance (Dobler et al. 2011; Santos et al. 2016). However, in cases where the cost of evolving additional autoresistance is low, the evolution of toxicity and resistance can become uncoupled.

Poison frogs of the family Dendrobatidae are a promising system to study the evolution of toxicity and autoresistance. The ability to sequester defensive alkaloids from dietary sources has evolved independently multiple times in this group (Santos et al. 2003; Vences et al. 2003; Santos and Cannatella 2011), and recent studies have identified amino acid substitutions on ion-transport proteins targeted by these toxins that coincide phylogenetically with the origins of alkaloid sequestration (Tarvin et al. 2016, 2017a; Yuan and Wang 2018). Some of these changes have been shown to provide toxin resistance *in vitro* (Tarvin et al. 2017a; Wang and Wang 2017).

Within Dendrobatidae, the genus *Phyllobates* is unique for secreting Batrachotoxin (BTX; Märki and Witkop 1963; Myers et al. 1978), one of the most powerful neurotoxins known to science (LD_50_ = 2μg/kg subcutaneous in mice; Tokuyama et al. 1968). Although several poison frog species from other genera (e.g. *Andinobates*, *Dendrobates*, *Oophagaa*) coexist with *Phyllobates* (Silverstone 1976; Myers et al. 1978), and feed on relatively similar prey types (Toft 1981; Caldwell 1996; Arce and Rengifo 2013; Osorio et al. 2015), decades of chemical work on skin extracts from more than 70 species of poison frogs (Daly 1998; Daly et al. 2005; Santos et al. 2016) have only found BTX on *Phyllobates* species. This steroidal alkaloid binds to the ***α*** subunit of voltage-gated sodium channels on nerve and muscle cells, reducing their affinity for Na^+^ ions, and leaving them permanently open and unable to experience action potentials (Märki and Witkop 1963; Daly et al. 1965; Warnick et al. 1976; Strichartz et al. 1987; Wang et al. 2006). Yet, nerve and muscle membranes of *Phyllobates aurotaenia* and *P. terribilis* are essentially insensitive to the action of BTX (Albuquerque et al. 1973; Daly et al. 1980). Even captive-bred individuals that were never exposed to BTX (which is obtained from dietary sources) showed full resistance, suggesting a strong genetic component of autoresistance (Daly et al. 1980). *Phyllobates* species vary widely in the amount of BTX stored in the skin, ranging from almost undetectable levels (∼0-1µg per frog) in *P. vittatus* and *P. lugubris* (Daly et al. 1987) to astoundingly high quantities (∼700-1900µg per frog) in *P. terribilis* (Myers et al. 1978). Furthermore, toxicity has increased at least twice in the evolutionary history of this genus, once along the branch leading to *P. aurotaenia*, *P. bicolor* and *P. terribilis*, and again in the lineage that gave rise to *P. terribilis* (Fig. 1; Myers et al. 1978; Daly et al. 1980, 1987), making this genus a fitting system to study the evolution of autoresistance.

**Figure 1.**
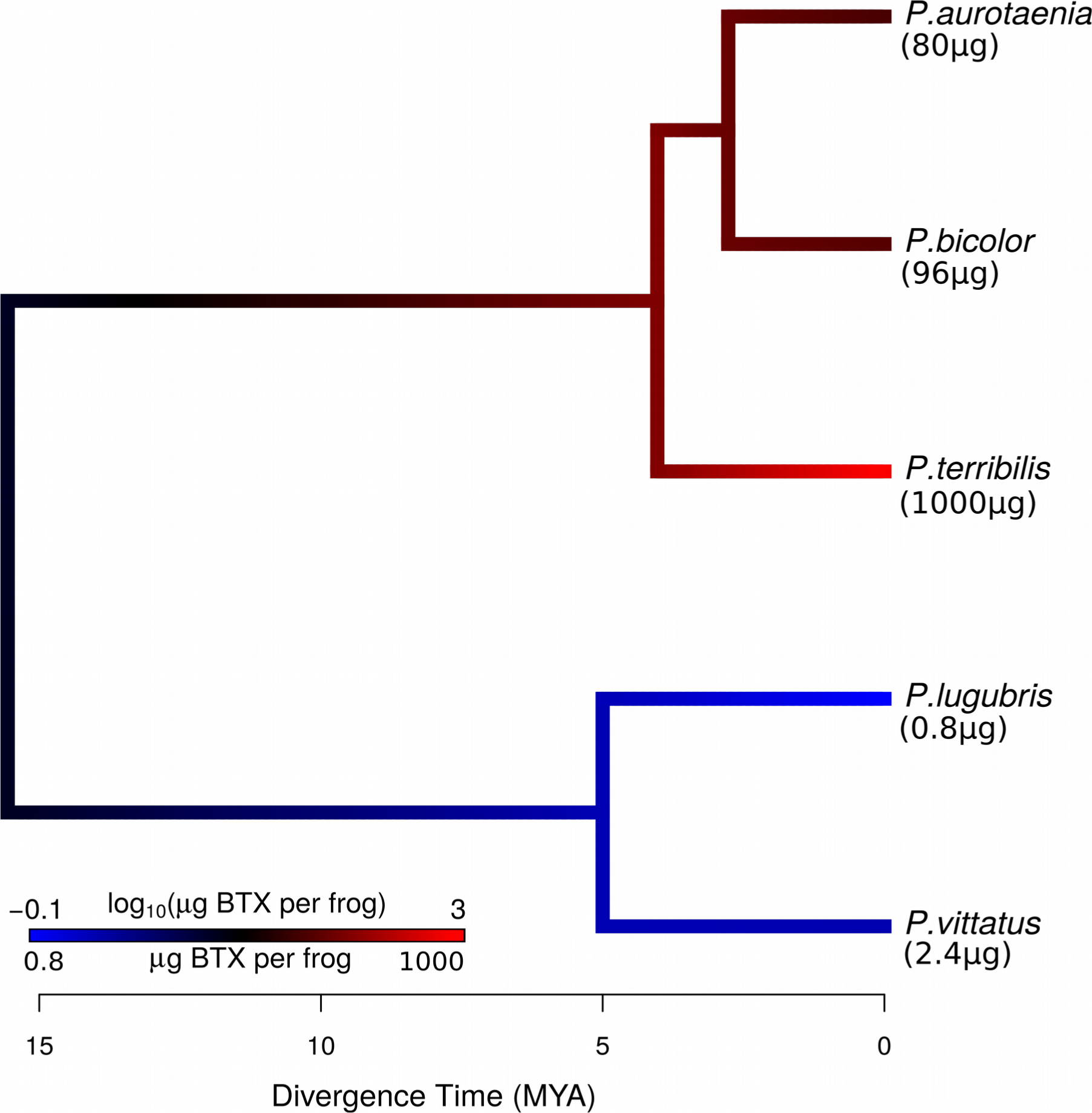
Phylogenetic relationships and average levels of cutaneous BTX per frog among *Phyllobates* species, estimated from pooled batches of frog skins, as presented in Table 2 of Daly et al. (1987). Branches were colored based on a maximum likelihood ancestral state reconstruction under Brownian Motion using the approach of Revell (2013; Method 2). The topology follows Grant et al. (2017), and divergence times were obtained from the TimeTree portal (Kumar et al. 2017). Numbers above the color bar are in log_10_ units, whereas those below the bar are in standard units.

Tarvin et. al. (2016) identified five amino acid replacements (A423S, I433V, A446D, V1583I, N1584T; numbering follows positions on the rat sequence) at or close to sites known to interact with BTX on the S6 segments of domains DI and IV of the muscular voltage-gated sodium channel (Na_V_ 1.4, encoded by the SCN4A gene) of *P. terribilis.* One of them (V1583I) was also present in *P. aurotaenia*. Further work (Wang and Wang 2017) showed that only N1584T provides BTX resistance *in vitro* when introduced onto the rat Na_V_ 1.4. Multiple combinations of the five substitutions were tested, and only those where N1584T was present (including N1584T alone) conferred BTX resistance.

Based on the data available from these two species, autoresistance seems to have evolved in tandem with increases in BTX levels, with *P. terribilis* having accumulated more mutations at BTX-sensing residues and greater BTX resistance than the less toxic *P. aurotaenia*. However, previous electrophysiological experiments have shown that nerve and muscle fibers of both *P. terribilis* and *P. aurotaenia* remain fully functional in the presence of BTX concentrations that completely inactivate the same tissues in other frogs, namely *Rana pipiens* (Albuquerque et al. 1973) and the dendrobatid *Oophaga histrionica* (cited as unpublished in Daly et al. 1980), indicating that both *Phyllobates* species are highly resistant to BTX. Furthermore, it was recently suggested that some of the amino acid differences observed between *P. terribilis* and *P. aurotaenia* could due to sequencing artifacts (Yuan and Wang 2018), so the extent to which the SCN4A genotypes of these two species differ is unclear.

Our aim here is to further elucidate the history of autoresistance-related mutations in *Phyllobates* SCN4A genes, in order to evaluate the extent to which BTX autoresistance has coevolved with toxicity levels in this group. To do so, we have generated SCN4A sequences from all known species of *Phyllobates,* representing the broad spectrum of BTX variation in this group (Fig. 1), which allows us to test this correlation beyond *P. terribilis* and *aurotaenia*. If autoresistance is indeed correlated with BTX levels, species with higher BTX contents should exhibit more resistant genotypes (e.g. with a higher number of AA changes at BTX sensing sites).

## Methods

Combining data from previous work (Tarvin et al. 2016; Yuan and Wang 2018) and newly generated sequences, we amassed a dataset of SCN4A sequences from 147 individuals of 45 species (35 Dendrobatoids and 10 outgroups; Tables 1 and S1), including the five known species of *Phyllobates* (36 samples; 2-14 per species). Alkaloid profiles are available for 30 of the 35 dendrobatoids used (Table 1), which allows us to confidently assume that, at least among the species sequenced BTX secretion originated at the base of *Phyllobates*. Our dataset encompasses the S6 and P-loop segments of Domains I-IV of the SCN4A gene. These regions are located on the pore of the Na_V_1.4 channel, where BTX binds, and *in vitro* directed mutagenesis studies have uncovered over a dozen mutations that confer BTX resistance to mamalian and insect voltage-gated sodium channels at these sites (Table S2). Furthermore dendrobatid frogs, including *Phyllobates* have mutations related to autoresistance at some of these regions (Tarvin et al. 2016). We then used ancestral sequence reconstructions to investigate the evolutionary history of these segments in relation to the acquisition and further increases in BTX-based toxicity among *Phyllobates* species.

**Table 1.**
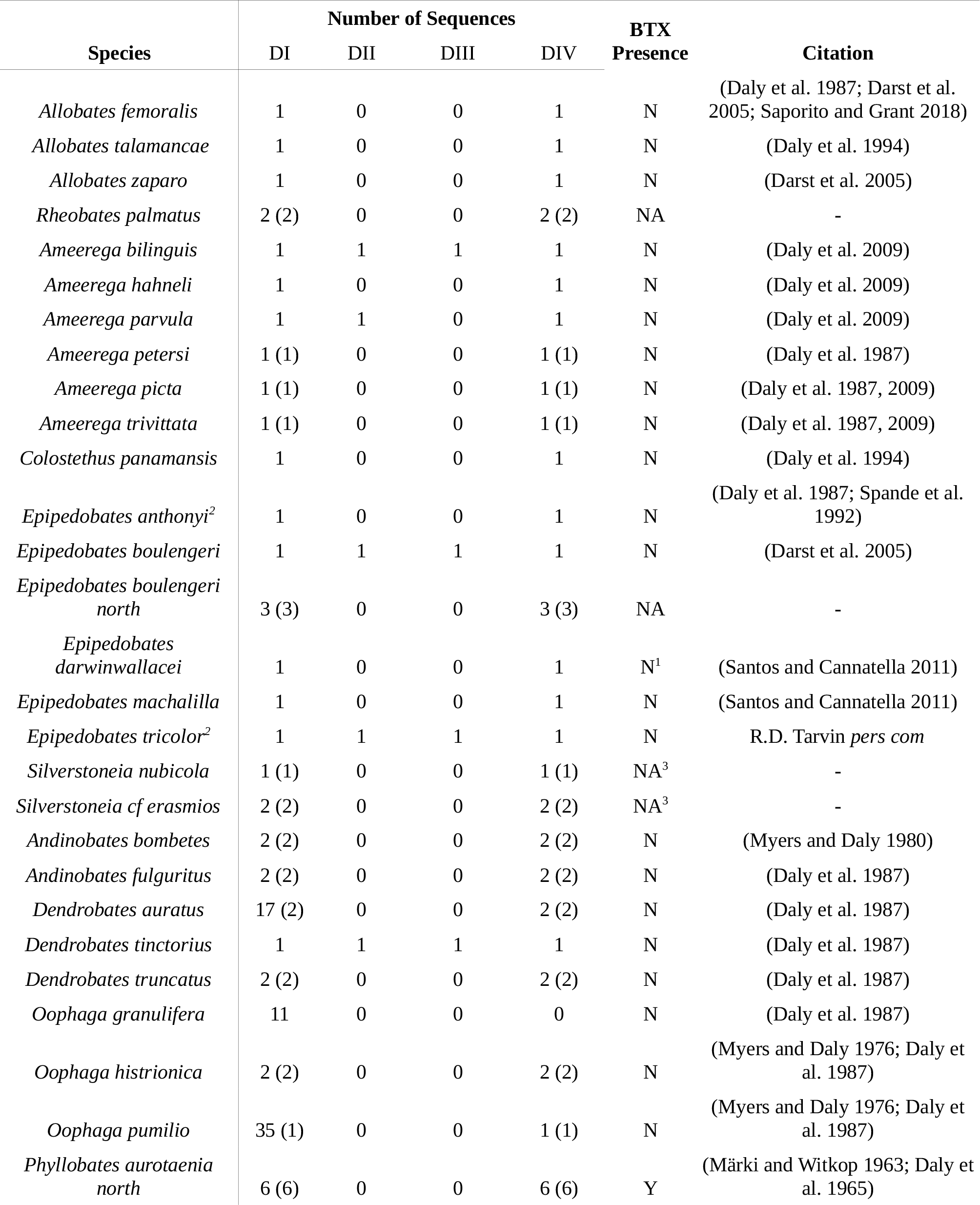
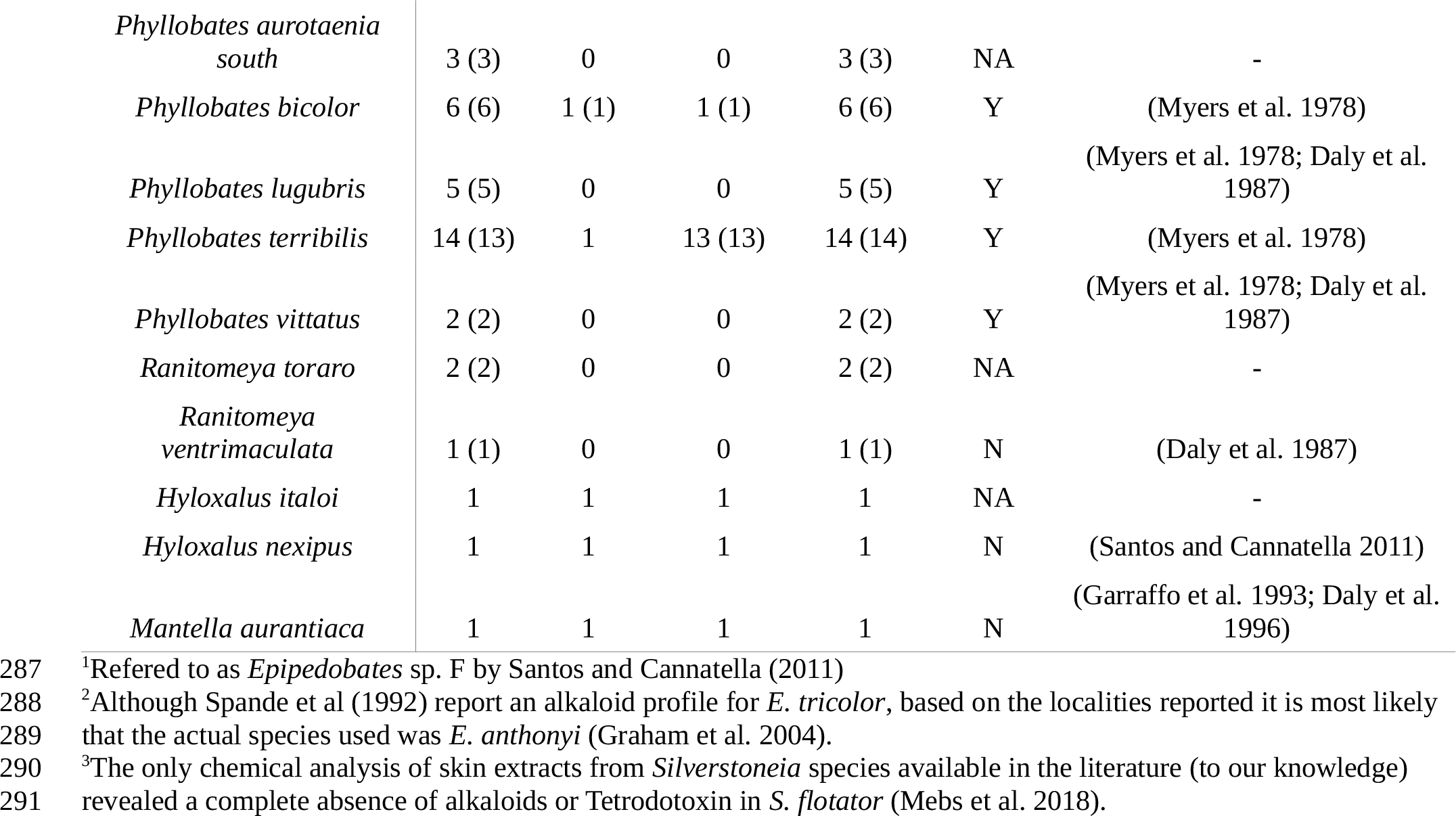
Number of sequences analyzed per SCN4A segment and presence/amount of BTX in the species of poison frogs used in this study. Numbers in parentheses represent new sequences added for this study.

### Publicly available data

We downloaded publicly available SCN4A sequences for 27 dendrobatid species and seven outgroup anuran species (Table S1). Sequences from seven specimens were excluded following previously outlined concerns (Yuan and Wang 2018; Table S3). In addition, we extracted SCN4A sequences from recently published transcriptomes of *Rana pipiens* (http://www.davislab.net/rana/; Christenson et al. 2014) and *Rhinella marina* (http://gigadb.org/dataset/100374; Richardson et al. 2017), and the genome of *Nanorana pariekri (V2, http://gigadb.org/dataset/100132; Sun et al. 2015)*. To do so, we querried the *Xenopus tropicalis* Na_V_ 1.4 protein sequence (ENSXTP00000031166) against each transcriptome/genome annotation with tblastn, and retained the best hit. We then confirmed orthology of these sequences to SCN4A using the phylogenetic approach detailed in the *Sequence analysis* section below.

### DNA sequencing

We sequenced the S6 segments of the DI and DIV domains from 59 individuals of 20 species of dendrobatids, 12 of which were not previously represented in public databases. DNA was extracted from toe-clip, mouth swab or liver samples using Qiagen DNeasy spin columns, and SCN4A fragments were amplified with primers designed based on the *X. tropicalis* sequence (ENSXETG00000014235), and refined as new sequences were generated. Table S4 contains primer sequences and thermal cycling protocols. PCR products were purified with ExoSap and Sanger-sequenced in both directions to confirm base calls.

### Transcriptome sequencing

We obtained a full SCN4A mRNA sequence for *P. bicolor* from a transcriptome assembly generated for an ongoing project (Márquez, R. et al. unpublished). RNA was extracted from skin, liver, heart, and muscle tissue, pooled in equimolar ratios, used to build a paired-end cDNA library, and sequenced on an Illumina HiSeq 2000. After quality trimming and adapter contamination removal with Trimmomatic (Bolger et al. 2014), we used Trinity (Grabherr et al. 2011) to generate an assembly. We then obtained the SCN4A sequence as described above for other species.

### Sequence analysis

For each of the four SCN4A fragments, we aligned all homologous sequences of each species to extract unique haplotypes, which were then aligned across species. From these alignments we built maximum likelihood trees for each segment to search for possibly contaminated sequences (Fig. S1). Next, one protein sequence was randomly selected per species for further analyses, except for *P. terribilis*, where two alleles had amino acid differences in the DI-S6 segment, so we kept both alleles. Finally, to confirm orthology of the protein sequences in our dataset (including those derived from genomes and transcriptomes) to SCN4A, we aligned them to sequences of all genes in the SCNA family from other vertebrates available in ENSEMBL and built a maximum likelihood tree (Fig. S3). All alignments were done using MUSCLE (Edgar 2004), and all trees were built using PhyML (Guindon and Gascuel 2003; Guindon et al. 2010) under sequence evolution models chosen with ProtTest (Darriba et al. 2011) or jModelTest (Darriba et al. 2012).

In order to infer the phylogenetic origin of amino acid substitutions, we conducted ancestral sequence reconstructions in PAML (Yang 2007). Each SCN4A fragment was analyzed independently under the best protein evolution model selected by ProtTest. We provided PAML with a topology based on Grant et al. (2017) for dendrobatoid relationships and Pyron and Wiens (2011) for outgroup relationships (Figs 2-3), and optimized its branch lengths during each ancestral reconstruction. Some populations of *Phyllobates aurotaenia* and *Epipedobates boluengeri* present in our dataset have been suggested to be distinct non-sister lineages by recent studies (Grant et al. 2017; Tarvin et al. 2017b), so we represented them as such in our phylogenies.

**Figure 2.**
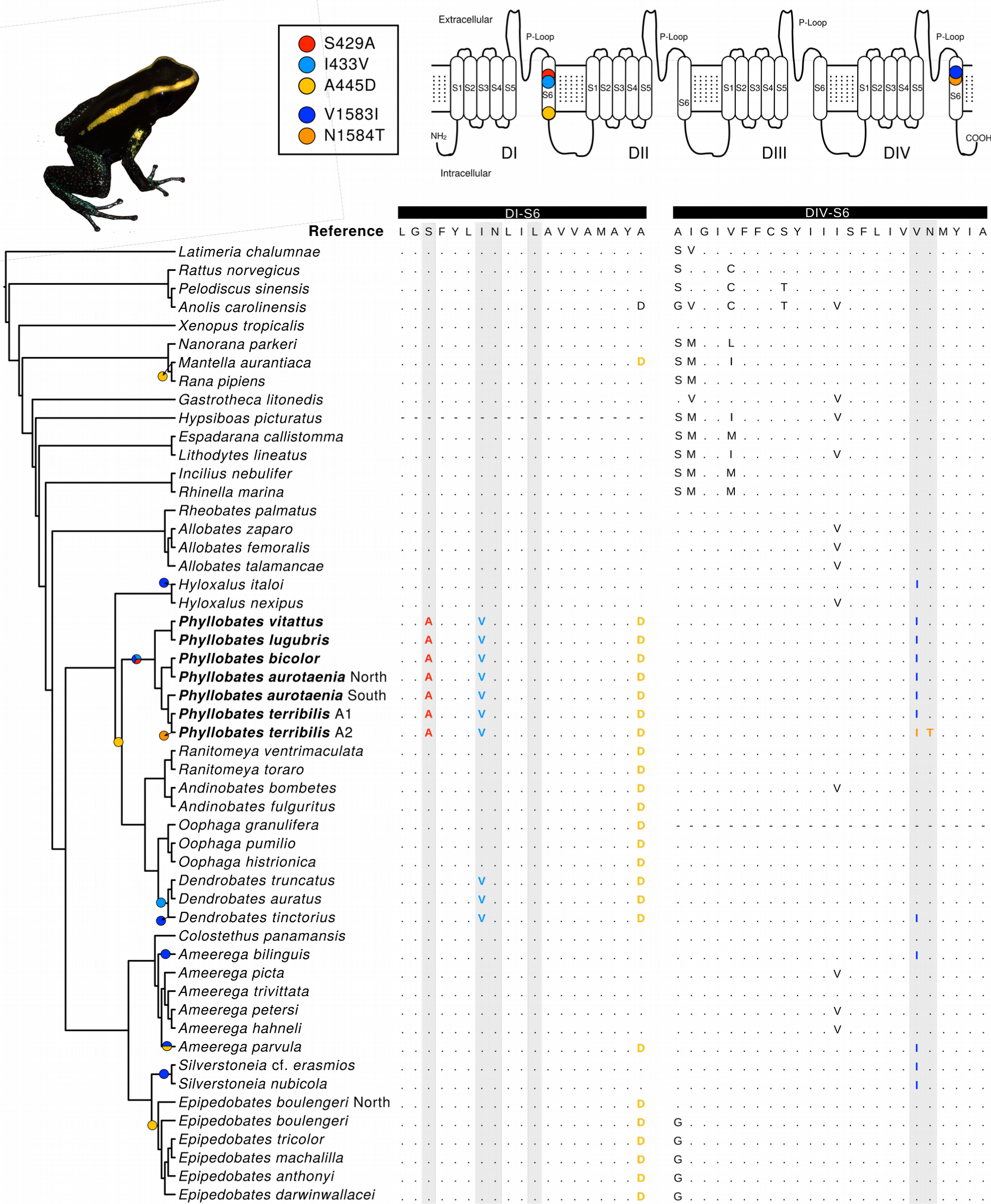
Amino acid sequences and ancestral reconstructions of the DI and DIV S6 segments of dendrobatids and other frogs. The reference sequence corresponds to the reconstructed ancestral frog sequence. The location of substitutions potentially important for autoresistance is indicated on the Na_V_1.4 schematic above the alignment, and the origin of each substitution is indicated on the corresponding branch. Sites known to be involved in BTX binding (Table S4) are shaded in grey. The topology follows Grant et al. (2017) and Pyron and Wiens (2011), and branch lengths are not meaningful. Non-anuran sequences (i.e. coelacanth, rat, turtle, and anole) are only shown for comparison, and were not used in analyses.

## Results

In concordance with previous work (Tarvin et al. 2016), we found five substitutions on the S6 segments of domains DI (S429A, I433V, A445D) and DIV (V1583I, N1584T) in *Phyllobates* frogs (Fig. 1). According to our ancestral sequence reconstructions, three of them (S429A, I433V, V1583I) arose at the root of the genus *Phyllobates,* coinciding with the acquisition of BTX secretion. A445D evolved earlier, at the common ancestor of *Phyllobates*, *Dendrobates*, *Ranitomeya*, *Andinobates*, and *Oophaga* (i.e. the subfamily Dendrobatinae *sensu* Grant et al. 2006, 2017). Surprisingly, N1584T, the only substitution shown to confer BTX resistance on rat Na_V_1.4 channels (Wang & Wang, 2017), was present only in a single individual of the 14 *P. terribilis* sequenced, and not found in any other species.

We did not find any substitutions coinciding with the origin of BTX secretion in other regions of SCN4A known to interact with BTX (i.e. DII and DIII S6 segments and DI-IV P-loops; Wang et al. 2000; Wang et al. 2001; Wang et al. 2006; Fig. 2, Fig. S2). However, our ancestral reconstructions uncovered five previously unreported substitutions in these regions (Y383F, F390Y, V748I, V774T, M777L; Fig. 3) that originated in alkaloid-sequestering clades, including the ancestor of Dendrobatinae (Y383F, F390Y, V774T, M777L). Our reconstructions show that four of these substitutions (Y383F, V748I, V774T, M777L) evolved more than once (although note that, despite high posterior probabilities [all > 0.95], taxon sampling for DII and DIII is sparse), and Y383F and M777L are present in *Mantella aurantiaca*, a member of a separate, distantly related radiation of poison frogs that convergently evolved the ability to sequester many of the same alkaloids present in dendrobatids (Garraffo et al. 1993; Daly et al. 1996). Furthermore substitution M777L was found to have occurred in parallel in five of the six frog SCNA paralogs in the recent history of dendrobatids (represented by *Oophaga pumilio;* Rogers et al. 2018). These results suggest a potential role of these five substitutions in alkaloid autoresistance that deserves further investigation.

**Figure 3.**
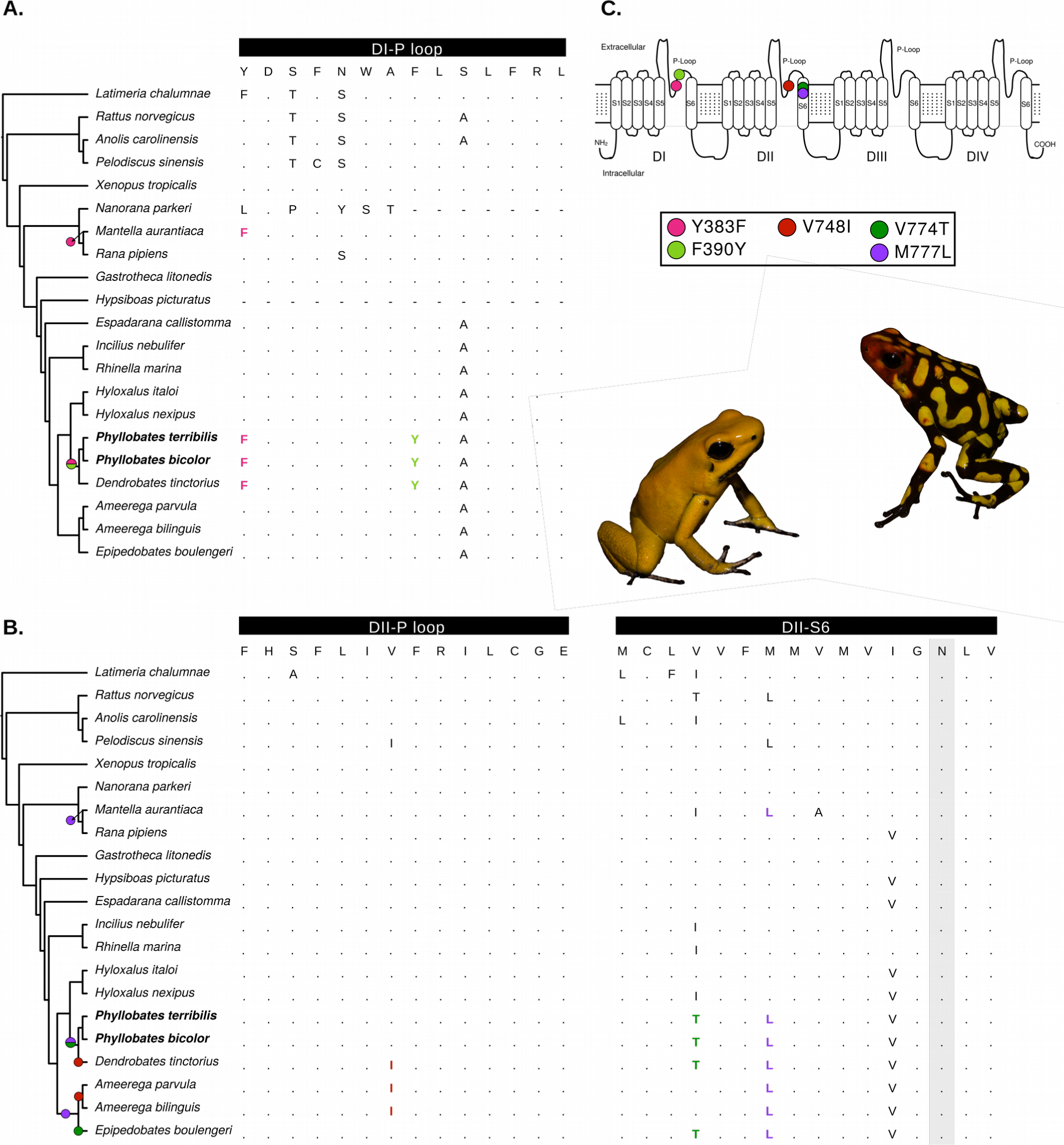
Amino acid sequences and ancestral reconstructions of the DI P-loop (A) and the DII P-loop and S6 segments (B). The locations and evolutionary origins of mutations are shown in panel C and on the phylogeny. The topology and shading are as in Figure 2.

## Discussion

Ancestral sequence reconstructions show that most of the amino acid substitutions in BTX-sensing regions of *Phyllobates* SCN4A alleles either predated or coincided with the evolution of BTX secretion, with the exception of N1584T, which seems to have evolved recently within *P. terribilis*, where it is still polymorphic and at low frequency. This points to a scenario where the ancestral *Phyllobates* Na_V_1.4 protein acquired autoresistance in concert with the evolution of basal levels of BTX, possibly facilitated by preexisting substitutions, and did not evolve further autoresistance as toxicity increased. In other words, our results suggest that the ancestral lineage of *Phyllobates* evolved sufficient BTX resistance at the Na_V_1.4 channel to withstand the broad range of toxicities currently present in its descendants. Once this high basal level of BTX resistance was acquired, the evolution of increased toxicity was released from the costs of increasing autoresistance.

This conflicts with the functional work of Wang and Wang (2017), who showed that neither of the three mutations coinciding with BTX secretion (S429A, I433V, V1583I; Fig. 2) nor their combinations decrease the susceptibility of the rat Na_V_1.4 to BTX. However, to fully understand the functional effects of these mutations on poison frog channels it is necessary to consider the genetic background on which they arose, since differences between the rat and *Phyllobates* channels at other sites, such as those identified in the DI P-loop and DIIS6 (Fig. 3), are likely to influence the interactions of sites 429, 433, and 1581 with BTX. For example, Tarvin et al. (2017) recently found an important effect of the genetic background (poison frog vs. human) when performing site-directed mutagenesis tests of epibatidine resistance. Although substitutions S429A, I433V and V1583I could presumably be involved in resistance to other alkaloids, they all occur at sites of demonstrated relevance in BTX binding to Na_V_1.4 (Wang and Wang 1998; Vendantham and Cannon 2000) or Na_V_1.5 (Wang et al. 2007) channels, and mutations at these sites (different from those in *Phyllobates*) confer BTX resistance to mammalian channels *in vitro* (Wang and Wang 1998; Vendantham and Cannon 2000; Wang et al. 2007; Table S2), suggesting an important role in the evolution of BTX autoresistance. Furthermore, although none of the mutations that predated the acquisition of BTX are on sites known to interact with BTX, many are close to these sites, which leads us to suspect that at least some of them may have influenced the evolution of BTX autoresistance, and could therefore explain the discordance with the results of Wang and Wang (2017). Biochemical assays that examine the effect of mutating *Phyllobates* sequences back to ancestral genotypes should provide insight on the functional and evolutionary implications of specific mutations in BTX autoresistance. In the meantime, the molecular and physiological mechanisms behind BTX resistance in *Phyllobates* remain an open question.

It is possible that N1584T played a role in the evolution of resistance to the very high levels BTX found in *P. terribilis*, since this and other mutations at this residue confer BTX resistance to rat Na_V_1.4 channels *in vitro* (Wang and Wang 1999, 2017). However, the fact that this mutation occurs at low frequency in *P. terribilis* lends little support to this hypothesis. Even the lowest amount of cutaneous BTX observed in individuals of this species (∼700μg) is much higher than those found in any other species (Myers et al. 1978; Daly et al. 1980, 1987). Had N1584T played an important role in allowing this increase in toxicity we would expect it to have rapidly become fixed by positive selection. Further investigation of allele frequencies at this site and BTX content variation in natural populations of *P. terribilis* could help clarify the role of N1584T in the evolution of BTX autoresistance.

Evolving neurotoxin-resistant ion channels many times involves mutations at functionally important residues, which are therefore likely to have negative pleiotropic effects. For example, several mutations that make sodium channels resistant to Tetrodotoxin (TTX), a Na_V_ blocker, have been shown to negatively impact the channel’s voltage-gating and permeability/selectivity properties (Chiamvimonvat et al. 1996; Pérez-García et al. 1996; Lee et al. 2011). In addition, substitutions that provide resistance to higher concentrations of TTX also tend to produce greater reductions in channel performance (Feldman et al. 2012). Therefore, populations of the TTX-resistant snake *Thamnophys sirtalis* appear to fine-tune their degree of TTX resistance based on the toxicity of their local newt prey (Brodie et al. 2002).

All known BTX-resistant mutations (Table S2) are located on or close to sites crucial to channel function, such as the gating hinge (formed by residues G428, G783, G1275, S1578; Zhao et al. 2004), or the ion selectivity filter (i.e. the DEKA locus; residues D400, E755, K1237, A1529; Backx et al. 1992; Favre et al. 1996), which could promote a similar correlation in BTX-resistant *Phyllobates* sodium channels. Our results, nonetheless, provide no evidence in favor of this scenario. This could be due to several reasons. For example, it is possible that increased resistance has evolved in more toxic lineages via alternative mechanisms, such as toxin modification or sectorization. The fact that isolated nerves and muscles of *P. terribilis* and *P. aurotaenia* resist high levels of BTX (Albuquerque et al. 1973; Daly et al. 1980), however, makes this an unlikely scenario. Another (non-exclusive) possibility is that the combination of mutations S429A, I433V, and V1583I may provide high BTX resistance at a low functional cost, and that this genotype arose through an accessible mutational pathway, reducing the extent of selection against highly autoresistant genotypes in low-toxicity individuals. Additional studies that address the functional effects of these mutations in terms of BTX resistance and sodium channel/muscle performance should disentangle this issue.

Finally, our data also contribute to the understanding of the general patterns of autoresistance evolution in poison frogs, which accumulate many different toxic alkaloids. We inferred several mutations evolving at the roots of alkaloid-defended clades (e.g. Y383F, 445D, V774T, V777L), while others appear later within these clades, in closely related species with similar alkaloid profiles (e.g. S429A, V433I). This pattern is compatible with a scenario involving initial adaptation to a basal toxin profile followed by (and possibly allowing for) further diversification and increased complexity of chemical defense (e.g. more diverse alkaloid profiles) among toxic clades (Santos et al. 2016; Tarvin et al. 2016). Many of these changes occurred in parallel between alkaloid-bearing lineages, even dendrobatids and mantellids, which diverged ∼150 MYA (Kumar et al. 2017). Such parallelisms may be due to strong functional constraints on sodium channel evolution, although other explanations such as historical contingency can not be discarded (Wright 1932; Dean and Thornton 2007; Stern and Orgogozo 2009).

Overall, our results suggest that *Phyllobates* poison frogs evolved BTX-resistant Na_V_1.4 sodium channels in concert with the ability to secrete this toxin, and that the basal level of BTX resistance was high enough to support toxicity increases throughout the evolution of the genus without evolving further autoresistance. Future studies integrating biochemistry, physiology and population genetics are needed to illuminate the functional and evolutionary mechanisms driving the evolution of BTX resistance in these frogs’ sodium channels, especially the functional effects of SCN4A mutations in relation to the tradeoff (or absence thereof) between BTX resistance and sodium channel performance.

## Acknowledgements

We thank Chris R. Feldman, Jorge A. Molina, Andrew J. Crawford, and Mohammad A. Siddiq for insightful comments and suggestions, Valentina Gómez-Bahamón and Daniel R. Matute for comments on the manuscript, Ralph A. Saporito and for clarifications about poison frog alkaloid profiles, Rebecca D. Tarvin for sharing unpublished results, and Álvaro Hernandez for assistance with transcriptome sequencing. This work was supported by a Seed Grant from the Faculty of Sciences at Universidad de los Andes to R.M., a Basic Sciences Grant from the Vicerectory of Research of the same institution to A.A. and R.M., and NSF Doctoral Dissertation Improvement Grant (DEB-1702014) to R.M. and Marcus R. Kronforst. R.M. was partially supported by a Fellowship for Young Researchers and Innovators (Otto de Greiff) from COLCIENCIAS. Tissue collections were authorized by permits No. 004, 36, 2194, and 1177 granted by the Colombian Ministry of Environment and Authority for Environmental Licenses (ANLA), 078-2003 from the Peruvian National Insititute of Natural Resources, 002-012012 from the Nicaraguan Environment and Natural Resource Ministry (MARENA), and SC/A-37-11 from the Panamanian Genetic Resource Access Unit (UNARGEN).

